# Epidemic size and host–parasite (co)-evolution: a meta-analysis

**DOI:** 10.64898/2026.09.18.752591

**Authors:** Sam Paplauskas

**Affiliations:** University of Stirling, Department of Biological and Environmental Sciences, Stirling, FK9 4LA, Scotland, UK

## Abstract

The size of an epidemic can influence the strength of selection acting on hosts and parasites, but whether epidemic size consistently shapes host–parasite (co)-evolution remains unclear. I conducted a meta-analysis of studies that simultaneously quantified epidemic size and evolutionary change in hosts, parasites or their interactions, synthesising 123 effect sizes from 11 studies. I tested whether epidemic size was associated with four components of evolutionary change: host evolution, parasite evolution, non-additive host–parasite coevolution and net coevolution. Larger epidemics were associated with greater host evolutionary change in invertebrates, whereas the relationship was positive in plants and negative but non-significant in bacteria. Parasite evolution, non-additive coevolution and net coevolution were not detectably associated with epidemic size, although parasite infectivity showed a relatively consistent increase across studies. These results provide limited evidence for a general relationship between epidemic size and host–parasite (co)-evolution and instead suggest that evolutionary responses depend on the host–parasite system and broader ecological and temporal context. Interpretation is constrained by the small and taxonomically biased evidence base, particularly the strong representation of *Daphnia*, and by substantial heterogeneity in how epidemic size and evolutionary change were measured and defined. More studies across diverse host–parasite systems are required that track both evolutionary change and disease prevalence through time. This work also provides an empirical basis for incorporating coevolutionary feedbacks into quantitative epidemiological models.

## Introduction

Epidemics are periods during which the incidence or prevalence of a disease increases substantially within a host population, resulting in elevated exposure to infection and increased disease burden (Jenelle et al., 2007). By reducing host survival, reproduction and population abundance, epidemics can have substantial ecological consequences for host populations (Altizer, 2003). Studies of host–parasite interactions suggest that these effects can also create opportunities for reciprocal selection: infection may favour host traits that reduce susceptibility or the fitness costs of infection, while parasites may be selected for traits that increase transmission or overcome host defences (Koskella and Lively, 2009; Paterson et al., 2010; Schulte et al., 2011). Such reciprocal selection provides the basis for evolutionary change in hosts and parasites and, where evolutionary change in one population alters selection on the other, for host–parasite coevolution (Paplauskas et al., 2021). However, whether the magnitude of an epidemic is actually associated with the extent of such evolutionary change is not well-established.

Based on the general principles of natural selection, we might expect the magnitude of an epidemic to influence the extent of evolutionary change in host and parasite populations. Larger epidemics involve a greater proportion of infected individuals within the host population (Dicker, 2006), and therefore expose more hosts to parasite-mediated selection. Where hosts differ genetically in their susceptibility or tolerance to infection, this greater exposure could produce a greater change in the frequencies of host genotypes across generations and, consequently, greater evolution of infection-related characteristics. Here, evolution refers to a change in the frequencies of heritable traits or alleles within a population over generations, rather than the generation of new mutations. Mutation generates new genetic variation on which selection can act, whereas evolutionary change describes changes in the representation of existing genetic variation within a population. In practice, such evolutionary change may be measured as changes in mean host infection phenotypes between generations (Brockhurst & Koskella, 2013).

The relationship between epidemic size and parasite evolution may be less straightforward. A parasite’s ability to infect and reproduce within a host is expected to be under strong selection because failure to infect can carry substantial fitness costs, including loss of reproduction or mortality (Salathé et al., 2008). If this selection operates whenever parasites encounter hosts, its strength may not necessarily increase in direct proportion to epidemic size. Nevertheless, the evolutionary responses of both hosts and parasites will depend on multiple interacting factors, including the costs of resistance, the strength of host- and parasite-mediated selection relative to other biotic and abiotic sources of selection (e.g. temperature, productivity and predation; Boots et al., 2009; Duffy & Forde, 2009; Koskella, 2018), and asymmetries in the evolutionary potential of host and parasite populations (Schmid-Hempel, 2011).

Theoretical studies further suggest that epidemiological dynamics can influence the strength of selection acting on hosts and parasites. In an early model incorporating evolutionary processes into epidemiological models, Anderson and May (1982) showed that host resistance was expected to evolve when parasite transmission was intermediate and resistance carried a fitness cost. At very high levels of transmission or virulence, however, parasite-induced mortality could cause host populations to decline, reducing the opportunity for selection to favour resistance. Similarly, models incorporating host–parasite coevolution and costs of resistance have predicted stronger selection for host resistance at intermediate levels of parasite virulence (Boots & Haraguchi, 1999). Models of parasite virulence and host resistance have further demonstrated that epidemiological feedbacks, including host mortality, can alter the strength of parasite-mediated selection (Gandon et al., 2001). Together, these studies suggest that epidemic dynamics have the potential to influence evolutionary responses, but do not establish how variation in epidemic size translates into variation in the extent of host or parasite evolution.

Empirical evidence directly linking epidemic size to the strength of selection and subsequent evolutionary change in host and parasite populations remains rare. Although evolutionary responses to parasite-mediated selection have been documented across a wide range of host–parasite systems, these studies have generally examined evolutionary change following infection without explicitly considering variation in the size of the epidemics generating that selection. For example, in human malaria, parasite-mediated selection has been implicated in shaping MHC diversity and the distribution of the sickle-cell trait (Hedrick, 2011; Kwiatkowski, 2005; Prugnolle et al., 2005). Similarly, amphibian populations exposed to large outbreaks of chytrid fungus have shown evidence of selection for increased resistance, although these outbreaks have also caused substantial population declines (James et al., 2015; Savage & Zamudio, 2011). Such studies demonstrate that infection can be associated with evolutionary change, but do not directly test whether variation in epidemic size is associated with variation in the extent of evolutionary change.

The few studies that have directly examined the relationship between epidemic size and evolutionary change have largely focused on *Daphnia*–microparasite systems. Duffy et al. (2012) found a positive relationship between epidemic size and the evolution of host resistance across seven *Daphnia dentifera–Metschnikowia bicuspidata* lake populations, suggesting that ecological gradients affecting epidemic size can influence evolutionary outcomes. However, the relationship was sensitive to the exclusion of an outlying population. In an experimental study of 16 replicate *Daphnia magna–Pasteuria ramosa* populations, Auld and Brand (2017) found that larger epidemics were associated with greater changes in host susceptibility and parasite within-host growth, while parasite infectivity increased regardless of epidemic size. More recently, Walsman et al. (2023) found that host resistance evolution could be weak when the costs of resistance outweighed its benefits. However, this study was based on only eight populations, used a peak-prevalence threshold of >0.1 to define epidemics, potentially encompassing relatively modest outbreaks, and examined a system characterised by low parasite virulence and low standing genetic variation. Together, these studies provide evidence that epidemic size can be associated with evolutionary outcomes, but the direction and generality of these relationships remain uncertain.

The predominance of *Daphnia* in this literature reflects the experimental tractability of these systems: *Daphnia* can be readily maintained in laboratory populations and have short generation times, allowing host–parasite evolution to be examined over relatively short periods, including within a single epidemic (Paplauskas et al., 2021). Nevertheless, even these studies have been constrained by relatively small numbers of replicate populations, limiting statistical power and making it difficult to establish general relationships between epidemic size and evolutionary change. Experimental studies of coevolution in longer-lived hosts, including plants, have also been conducted over multiple years (Thrall et al., 2012), but remain relatively uncommon. It therefore remains unclear whether the patterns observed in *Daphnia* represent general relationships between epidemic size and host and parasite evolution across host–parasite systems.

To address this lack of replication and assess the extent to which relationships between epidemic size and evolutionary change are consistent across host–parasite systems, I conducted a meta-analysis of 11 studies examining host evolution, parasite evolution, and both additive and non-additive forms of host–parasite coevolution (collectively, ‘(co)-evolution’; Paplauskas, 2025). Although the meta-analysis incorporated studies from multiple host–parasite systems, the available literature remained strongly biased towards *Daphnia* host–parasite systems, limiting the extent to which these relationships could be assessed across independent biological systems. Assuming no confounding factors, such as trade-offs between host resistance and fecundity, I predicted that (i) larger epidemics would be associated with greater reductions in host susceptibility to infection, (ii) changes in parasite infectivity would be independent of epidemic size, and (iii) net evolutionary change in infectivity would likewise be independent of epidemic size, reflecting the consistently strong selection expected to favour parasite infectivity.

Across the studies included in the meta-analysis, host evolutionary responses and net coevolutionary change showed substantial variation in direction, whereas parasite infectivity showed a more consistent increase. I found that this variation in host evolutionary responses was partly associated with epidemic size, with larger epidemics driving greater reductions in host susceptibility among invertebrates but increased susceptibility among plants. In contrast, parasite infectivity was comparatively consistent across epidemic sizes, and neither non-additive nor net coevolutionary change was significantly associated with epidemic size. These findings provide partial support for my predictions, with the relationship between epidemic size and host evolution consistent with stronger selection imposed by larger epidemics, while the relative consistency of parasite evolution was consistent with strong selection favouring parasite infectivity. However, the persistent taxonomic bias towards *Daphnia* systems limits the extent to which these patterns can be considered general across host–parasite systems.

## Methods

### Summary of the methods

To quantify the general relationship between epidemic size and the strength of host and parasite-mediated selection across host-parasite systems, I performed a meta-analysis to investigate the relationship between epidemic size and different components of host-parasite coevolution. After collecting a total of 436 studies from the current literature and assessing their suitability to my investigation, I ended up calculating 123 values for my effect size metric of interest, the log response ratio (lnRR). These effect sizes corresponded to the natural logarithm of the proportional change in a population-level expression of disease, such as the proportion of infected individuals within a population (Pinf) or the parasite transmission rate (β), between two points across an epidemic (such as pre and post-epidemic). Crucially, these changes in the population-level expression of disease were analysed in the context of their host-parasite (co)-evolutionary drivers using a revised version of the formulae by Paplauskas et al. 2021 and Paplauskas, 2025:

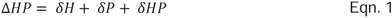

Where:

- ΔHP is the overall (i.e. ‘net’) change in a given metric of parasite infection success owing to additive host-parasite coevolution
- *δ*H is only the change in a given metric of parasite infection success owing to host evolution
- *δ*P is only the change in a given metric of parasite infection success owing to parasite evolution
- *δ*HP is only the change in a given metric of parasite infection success owing to non-additive host-parasite coevolution.

By splitting host-parasite coevolution into its constituent parts, I test the following three research questions (Q1-3):

Q1: What patterns of host-parasite (co)-evolution generally occur as a result of epidemics?

Q2: Does epidemic size determine the strength of host-parasite (co)-evolution?

Q3: What are some of the wider factors that influence host-parasite (co)-evolutionary responses to epidemics?

### Coevolution versus (co)-evolution terminology

Strictly speaking, host-parasite coevolution (in this study, also described as ‘net coevolution’) only refers to the sum of host evolution, parasite evolution and host-parasite non-additive coevolution (see equation 1). In comparison, host-parasite (co)-evolution offers flexibility when referring to a combination of one or more of host evolution, parasite evolution and / or host-parasite non-additive coevolution (Paplauskas, 2025). However, since it was only possible to calculate lnRR effect sizes for net host-parasite coevolution and two out of a possible total of three of its component parts (host evolution and parasite evolution cf. non-additive coevolution, see processing subsection of my data collection workflow) and untransformed values were used for my analysis of non-additive host-parasite coevolution (again, see processing), adhering to the strict definitions of host-parasite coevolution and host-parasite (co)-evolution could lead to misusing these terms interchangeably.

To avoid any confusion, rather than tediously separating ‘host-parasite coevolution’ and ‘host-parasite (co)-evolution’ terminology and to ensure consistent usage of ‘host-parasite (co)-evolution’ throughout, I extend my definition of host-parasite (co)-evolution to encompass any combination of host evolution, parasite evolution, non-additive host-parasite coevolution and net coevolution. This is particularly helpful for interpretation of i) the overall effect sizes for the orchard plot (includes host evolution, parasite evolution and net host-parasite coevolution) and ii) the full contextual factor analysis (includes host evolution, parasite evolution and net host-parasite coevolution).

### Data collection

Before collecting any data that would be used to calculate the log response ratio (lnRR) and a corresponding variance-covariance matrix of sampling errors, I developed a set of criteria that would be used to guide my search on the Google Scholar database for studies corresponding to my research question, and additionally would double as eligibility criteria during the second step of data collection (Table 2).

**Table 1.** Summary values for the set of two leave-one-out sensitivity analyses. Mean model coefficients ± the corresponding standard error are shown. These values were calculated by averaging across all meta-regression model coefficients generated by the iterative exclusion of either one study or independent treatment-control pairs (replicates) used when calculating effect sizes at a time. The following abbreviation is used; HType (Host type). Significant p-values (below a 95% significance threshold) are highlighted with bold font.

| Omission | (Co)-evol | HType | Estimate | Standard error | p-value |
| --- | --- | --- | --- | --- | --- |
| Study | Host | Invert. | -0.009 ± 0.001 | 0.002 ± 0 | **0 ± 0** |
| Study | Parasite | Invert. | 0.026 ± 0.021 | 0.01 ± 0.005 | 0.203 ± 0.177 |
| Study | Net | Invert. | 0.003 ± 0.001 | 0.006 ± 0.001 | 0.63 ± 0.136 |
| Replicate | Host | Invert. | -0.01 ± 0 | 0.002 ± 0 | **0 ± 0** |
| Replicate | Host | Plant | 0.004 ± 0 | 0.001 ± 0 | **0.02 ± 0.012** |
| Replicate | Host | Bacteria | -0.019 ± 0.001 | 0.01 ± 0.002 | 0.171 ± 0.043 |
| Replicate | Parasite | Invert. | 0.006 ± 0 | 0.004 ± 0 | 0.168 ± 0.024 |
| Replicate | Net | Invert. | 0.001 ± 0 | 0.006 ± 0 | 0.845 ± 0.012 |
| Replicate | Net | Plant | 0 ± 0 | 0.001 ± 0 | 0.751 ± 0.037 |

**Table 2.** Eligibility criteria.

| Criterion | Rationale |
| --- | --- |
| Any of metric of host-parasite (co)-evolution was measured (Equation 1), including host evolution ($\delta H$), parasite evolution ($\delta P$), net coevolution ($\Delta HP$) or non-additive coevolution ($\delta HP$). | Allowed me to test separate host, parasite and inter-dependent host-parasite evolutionary and coevolutionary hypotheses (as outlined in the introduction). Note that there was only one study with a measurement of non-additive coevolution (Paplauskas et al., 2021) for which the calculation of lnRR was not possible (see the fourth point of the *Data collection* section entitled *Processing*) so this was analysed using untransformed values. |
| The given metric of host-parasite (co)-evolution (referenced in (i)) was measured using a gold-standard approach to measuring (co)-evolutionary change, which was limited to studies including either a time-shift procedure that involved keeping one or both of the antagonists in evolutionary stasis (Brockhurst & Koskella, 2013) or a comparison of host genotype frequencies in ancestral and evolved populations (e.g. Duffy et al., 2008). | Strictly including only those studies with this gold-standard approach is intended to disambiguate between evolutionary versus coevolutionary change (Brockhurst & Koskella 2013; Paplauskas, 2025). It is worth noting that there were a limited number of studies combining both gold-standard approaches. |
| Epidemic size was measured using any | Where it had not been already calculated, |
| disease or infection-based metric of prevalence (the proportion of infected individuals within a population), or at least two instances of a given prevalence metric were measured across time. For studies involving experimental coevolution with bacterial hosts, the data necessary to calculate a proxy for epidemic size were measured (i.e. colony and plaque forming units per unit volume; Gomez & Buckling, 2011). | integrated epidemic size and its equivalent proxy for a single study of bacterial hosts (Gomez & Buckling, 2011) were calculated from the relevant host or parasite population data (described in the methods). Integrated epidemic size represents the time integral of the epidemic curve (i.e. area under the curve) and was my preferred choice to represent the selective pressure of an epidemic over one-off measurements of prevalence, such as peak prevalence, or the mean of multiple measurements. This was because these alternatives do not capture the temporal variation in prevalence like integrated epidemic size does, which not only incorporates multiple measurements of prevalence, but also incorporating information about the duration of the intervals between multiple measurements of prevalence. |
| A standard definition for what constitutes an epidemic was not required, such as a rapid increase in the proportion of infected individuals within a host population in a short period of time (Green et al., 2002). | This is because many studies vary in what defines an epidemic, and often there is no definition for what constitutes an epidemic (Paplauskas, 2025). |
| The data from the study was not already used by a different study included in the data I had collected as a re-analysis. | To avoid duplicating findings. |

After settling on the criteria outlined above (Table 2), the data collection process then involved four main steps (Fig 1):

1. *Identification*: I developed a combination of search terms (made up of keywords and phrases) according to my pre-determined set of eligibility criteria (Table 2) that were used to perform a literature search on the Google Scholar database on 7^th^ March 2023 (Table S1). This included a mixture of terms with variable scope that were included in five separate literature searches with default settings (records from all time, including citations) due to the search engine’s loose interpretation of complex nested Boolean Operators. These searches returned a total of 436 counts of matches potentially relevant for an investigation of the relationship between epidemic size and different components of host-parasite coevolution. Prior to screening, 15 records were lost during exportation due to an error syncing with the Google Scholar favourites library and two duplicate records were removed in R.
2. *Screening:* The first screening step involved the checking of titles and abstracts for relevance to the proposed research aim. Out of the 419 records involved at this first phase, 358 of them were excluded (224 and 134 titles and abstracts respectively). This was followed by downloading the full text of any remaining records (n = 61), reading them in detail and assessing them against the my pre-determined set of eligibility criteria (Table 2). Since the (co)-evolutionary data was not annotated by population id for one of the remaining records (Thrall et al., 2012), which prevented making the links to their epidemic data, this missing data was requested through personal communication with the lead author (whose reply was very much appreciated!).
3. *Extraction*: From the final 11 studies, the data necessary to calculate the effect size (lnRR) and its sampling variance were extracted. This included extracting the raw data for the mean, standard deviation and sample size for each comparison of host and parasite population-level phenotypes from the main article as text or figures (PlotDigitzer) and any supporting information files to calculate aggregate population-level summaries of the data or extracting these data directly from the published material (depending on availability). In addition, any data that could be used in accounting for the potential non-independence of replicates (i.e. ‘comparisons’) which were shared between the same study or experiment were recorded, along with standard meta-data (e.g. taxonomy), epidemic size and three other potentially key moderators of host-parasite (co)-evolution (see Table 3). It is important to note that many studies seemed to incorrectly label metrics of (co)-evolution (Paplauskas, 2025), which included the following:

- Auld and Brand, 2017: It is not possible to isolate the non-additive component of coevolution using their formula, but this was later updated in Paplauskas et al., 2021.
- Thrall et al., 2012: Referring to a comparison of past and future parasite infectivity against the corresponding contemporary host population for each timepoint is not ‘parasite evolution’ as described in their study, but it is in fact net coevolution.
- Gowler et al., 2023: Referring to a comparison of past and future host susceptibility against the corresponding contemporary parasite population for each timepoint is not ‘host evolution’ as described in their study, but it is in fact net coevolution.
- Strauss et al., 2017: According to Strauss et al. (2018), the susceptibility of host genotypes was measured using a set of infection experiments prior to the experimental coevolution. However, the parasite used to measure host susceptibilities was not frozen or otherwise prevented from evolving before carrying out the next stage of their experiment. Since the possibility of parasite evolution cannot be excluded, I decided that it was better to refer to ‘host evolution’ as net coevolution.
4. Processing: Finally, I processed the raw data extracted from six studies (Auld & Brand 2017; Duffy et al., 2008; Gowler et al., 2023; Strauss et al., 2017; Auld et al., 2014; Ameline et al.; 2022). This included calculating the population-level summary statistics, such as the mean, weighted mean (Strauss et al., 2017; Ameline et al., 2022), standard deviation and sample size that was not published in the original articles. Since some of the statistical information that could be used to calculate the standard deviation were missing for one particular study (Gomez & Buckling, 2011), these data were separately imputed for the pre and post-epidemic samples by averaging across the other values in my collected dataset (Furukawa et al., 2006). In addition, the mean values corresponding to the change in host phenotypic means for two studies was converted from resistance to susceptibility for consistency with the overall dataset (Gomez & Buckling, 2011; Ameline et al., 2022). This was achieved by simply using their complement (because susceptibility is the inverse of resistance) and, since it is a measure of spread, rather than location, the standard deviation did not require any adjustment. For all six of the studies in which the raw phenotypic data was processed, the variation in epidemic size over time was not already measured by the integrated epidemic size. Therefore, this was manually calculated as the area underneath the epidemic curve using the AUC function from the DescTools package in R (Signorell, 2024). The starting and finishing boundaries of the epidemic curve corresponded to when host and parasite phenotypic samples were taken for measuring different components of coevolution, rather than a standard epidemic definition (which was often missing, as discussed by Paplauskas, 2025). For the one study of a model bacteria-phage interaction (Gomez & Buckling, 2011), the conventional calculation for disease prevalence was adjusted to reflect the proportion of host bacteria by expressing infection (as plaque forming units) within the population (colony forming units plus plaque forming units). For the sole study in which the non-additive component of coevolution appeared (Paplauskas et al., 2021), it was not possible to convert the parasite transmission rate data into the appropriate format for calculating my effect size (see ‘Calculation of effect sizes (lnRR)’). This is because there were subzero mean values for the pre and post-epidemic transmission rate data, which caused an issue during log-transformation. Despite the possibility of retaining the data by ignoring the signs, the variable interpretation for different components of host-parasite coevolution would have led to possible confusion (the effect sizes for the non-additive component of host-parasite coevolution would have represented a ratio of magnitudes in comparison to a ratio of signed quantities for all others), so instead this transmission rate data was analysed using untransformed data.

**Figure 1.**
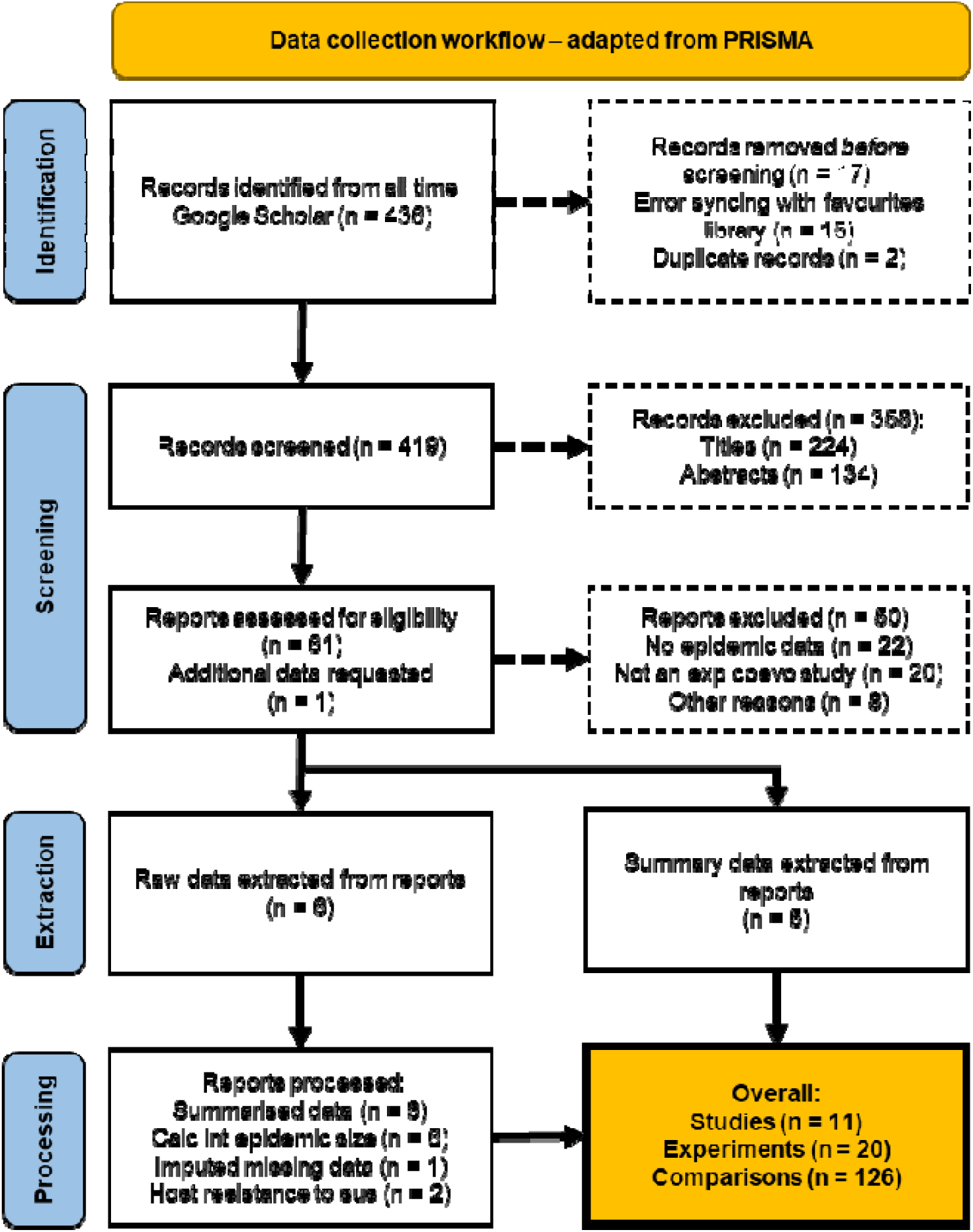
The four main steps of data collection. The final number of comparisons included in my dataset, which correspond to different components of host-parasite coevolutionary change in response to an epidemic, was 126 (and would be later used in the calculation of lnRR). Adapted from the PRIMSA workflow template (Page et al., 2021).

**Table 3.** Hypotheses describing how moderators (excluding epidemic size) included in this meta-analysis are predicted to influence host–parasite (co)-evolution during epidemics. To reflect the structure of the meta-analytical models, which simultaneously analysed host evolution, parasite evolution, and net coevolution as a single term, the term ‘host–parasite (co)-evolution’ is used here collectively to refer to these three components for which effect size data could be calculated.

| Moderator | Hypothesis |
| --- | --- |
| Metric of phenotypic change | As it is an unrealistic expectation, I suggest an alternative hypothesis to the underlying assumption that the different metrics of phenotypic change (i.e. (co)-evolution) can be treated as being |
| | equivalent to one another by the models used in this study. I predict that there is significant variation in host-parasite (co)-evolution during epidemics as a result of using different metrics of host-parasite (co)-evolution. This includes (i) the proportion of infected individuals within a host population ($P_{inf}$), (ii) the parasite transmission rate ($\beta$), (iii) the proportion of host clones within a population susceptible to parasite attachment ($P_{sus}$, as opposed to resistotypes *sensu* Ameline et al., 2022) and (iv) the proportion of plaque forming units on bacterial colonies (PFU, Gomez and Buckling, 2011). |
| Weighted mean | About half of the comparisons of phenotypic means were based on an unadjusted unweighted mean, whereas the other half were based on a frequency weighted mean approach. Since host individuals, genotypes and even resistotypes (Ameline et al., 2022) may vary not only in the magnitude and direction of (co)-evolution, but also by their frequency within a host population, this can lead to a potential bias in using unweighted data. Therefore, I predict that the range of error for host-parasite (co)-evolution during epidemics based on weighted means will be narrower than those based on unweighted means, and thus the overall weighted effect size is more likely significant. |
| Study setting | Similar to the expectation for the 'Weighted mean?' moderator, I predict that the range of error for host-parasite (co)-evolution during epidemics based on non-field studies will be narrower than those based on field studies. |

At the of the data collection (Fig. 1), there was a total 11 studies, 20 experiments and 126 comparisons that could be used to calculate effect sizes (lnRR).

### Calculation of effect sizes (lnRR)

Effect sizes were calculated as the log response ratio (lnRR). As mentioned in the summary of the methods, these effect sizes corresponded to the natural logarithm of the proportional change in average host-parasite infection phenotypes, such as the parasite transmission rate, between two epidemic timepoints (e.g. pre- and post-epidemic) relating to different components of host-parasite (co)-evolution (Equation 1; *sensu* Paplauskas et al., 2025). Each individual effects size (lnRR) was calculated using the following formula:

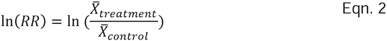

Where:

- *X̄_treatment_* = The mean population phenotype sampled at the end of the experiment, such as post-epidemic.
- *X̄_Control_* = The mean population phenotype sampled at the start of the experiment, such as pre-epidemic.
- Interpretation: The proportional change in the mean of the treatment relative to the control (e.g. the host population evolved 30% reduced susceptibility”).

Although there are different effect size metrics that could be used in this study, such as the standard mean difference (Borenstein et al., 2009; Field and Gillett, 2010), lnRR is a very popular metric for meta-analysis in ecology and evolution (Nakagawa et al, 2023) and is the the optimal metric for this study for the same reasons it is so popular. This is because the scale-free ratio of the log response ratio can be used to compare relative changes in data that originated on different scales. As a result, it is relatively easy to interpret versus other effect size metrics which can also compare data across scales, such as the standardised mean difference, which measures group differences in terms of standard deviations (Borenstein et al., 2009; Field and Gillett, 2010). The log transformation of the response ratio is also useful for interpretation of host-parasite (co)-evolutionary change data, as many of these trajectories can lead to negative values (e.g. host populations that evolve reduced susceptibility to infection), because it is symmetric around zero (in comparison to simply the untransformed response ratio, RR), but more importantly it has improved statistical properties for inference compared to simply the RR (Borenstein et al., 2009; Field and Gillett, 2010).

A matrix of sampling variances and covariances for lnRR was calculated using a standard formula (Borenstein et al., 2009; Field and Gillett, 2010). However, since there were no studies with multiple shared controls (or treatments), the final variance-covariance matrix was in fact equivalent to a vector of sampling variances. Since the approach of two studies in measuring pre-epidemic phenotypes led to a pair of unusually small pre-epidemic (control) phenotypes (equivalent to zero, Auld et al., 2014; Mitchell et al., 2004), a small value was added (0.01) to avoid inflating the final log response ratio during log transformation step. These studies were (i) a combined analysis of immune response and susceptibility in which there was no apparent evidence of host susceptibility to infection pre-epidemic (Auld et al., 2014) and (ii) a quantification of disease prevalence based on ‘the difference from a batch mean’ (Mitchell et al., 2004). Despite using a standard data handling approach, the effect size corresponding to an instance of parasite evolution was large relative to the rest of the data (lnRR = 4.14; Auld et al., 2014). Therefore, alongside the full dataset, a trimmed version of the dataset with this potential outlier removed was used in subsequent analysis.

### Mixed-effects meta-analytical models

A mixed effects meta-analytical model was fitted to the effect size data for lnRR that would form the basis for all of the future meta-analytical models used in this study. The same basic model structure was used in testing for publication bias and Q1-3 (summary of the methods). The mixed-effects meta-analytical model was developed using the rma.mv function from the metafor package v4.4.0 (Viechtbauer, 2010). This included (i) fixed effects for the log response ratio effect size and its variance-covariance matrix of sampling errors, (ii) standard random effects for study and both host and parasite genus, and (iii) nested random effects for each unique comparison from the same experiment. The standard random effects were used to account for the possibility of non-independence between experiments originating from the same study and the potential correlation between data collected from closely related host or parasite species. Similarly, correlated (nested) random effects were used to account for potential non-independence of comparisons taken from the same experiment (e.g. epidemics measured in multiple populations grouped by experimental treatments not accounted for in the moderator analysis).

### Publication bias

Before analysing the data fully, I used a funnel plot and Egger’s regression (Sutton, 2009) to test for the presence of publication bias. The test for publication bias was based on the aggregation of all three primary components of host-parasite coevolution focused on in this study, including host evolution, parasite evolution and net coevolution. Taking this approach to testing for the presence of publication bias ensured that the number of studies included the test exceeded the recommended threshold of 10 (Higgins, 2024). Although there are exceptions to this general rule of thumb (Mathes & Kuss, 2018), so the separate components of host-parasite coevolution could have examined independently, testing for the presence of publication bias is usually more effective when there are more than 10 studies included (Higgins, 2024). Also, it is worth mentioning that I adapted the mixed-effects model to be able to perform the Egger’s regression through the inclusion of the sampling variance of lnRR as a moderator.

Standard visual inspection of the funnel plot showed that there was no evidence of publication bias (Fig. S1). For instance, this was indicated by an approximately even distribution of effect sizes around the weighed average of the model. In support of this, there was no correlation between the size of the effects themselves and their standard error from the Egger’s test (R = -0.55, 95% CI [-1.11, 0.27], p = 0.43).

### Q1: General patterns of host-parasite (co)-evolution

Q1: What patterns of host-parasite (co)-evolution generally occur as a result of epidemics? This was quantified using two versions of the mixed-effects meta-analytical model (described earlier), including (i) a generic model that was fitted to the full dataset and (ii) another model that used a categorical moderator to account for any potential differences in how the different components of host-parasite coevolution might respond to epidemics. In addition, the sensitivity of the latter model to the inclusion of one extremely large effect size (lnRR = 4.14, Auld et al., 2014) was investigated by excluding it from the overall dataset and refitting the model to this reduced dataset.

As previously explained under the fourth point of the *Data collection* section (entitled *Processing*), only a single study measured non-additive host–parasite coevolution (Paplauskas et al., 2021), and its data could not be used to calculate lnRR. Therefore, to facilitate comparison with the mixed-effects meta-analytical model, the mean change in parasite transmission rate was instead compared with zero using a one-sample *t*-test, analogous to testing whether the mean individual effect size differed from zero.

The total variation in the standard mixed-effects model that included a categorical moderator was measured using the I-squared statistic. In addition, a breakdown of this relative heterogeneity was performed to measure how much each random effect level contributed to the relative heterogeneity.

### Q2: Size-specific patterns of host-parasite (co)-evolution

Q2: Does epidemic size determine the strength of host-parasite (co)-evolution? To address this question, whilst also simultaneously following-up on the heterogeneity analysis from Q1, a pair of meta-regression models (with and without the potential outlier described earlier) were developed from an adapted version of the mixed-effect analytical model (also described earlier) by specifying a three-way interaction term between the components of host-parasite (co)-evolution, host taxonomic classification and epidemic size (the latter of which was a continuous variable). The reason for including the host taxonomic classification within the interaction term, which broadly categorised hosts as invertebrate, plant or bacterial hosts, was to account for the inability to draw comparisons between the magnitude of host-parasite (co)-evolution where there is inconsistency in the scale on which host-parasite (co)-evolution is measured. This is because there is considerable variation between the generation time of invertebrate, plant and bacterial host species that affects their evolvability (Brockhurst & Koskella, 2013), and thus the rate at which evolution (or coevolution) can be observed for any given experimental design or unit of epidemic size.

As previously explained under the fourth point of the *Data collection* section (entitled *Processing*), only a single study measured non-additive host–parasite coevolution (Paplauskas et al., 2021), and its data could not be used to calculate lnRR effect sizes. Consequently, rather than conducting a meta-regression, the change in parasite transmission rate and epidemic size data were analysed using linear regression, allowing an analogous assessment of the relationship between epidemic size and non-additive host–parasite coevolution.

### Q3: Other context-specific effects of host-parasite (co)-evolution

Q3: What are some of the wider factors that influence host-parasite (co)-evolutionary responses to epidemics? Building on my other analyses of host-parasite (co)-evolution and epidemic size (Q1 and Q2), I investigated some of the general factors that could influence host-parasite (co)-evolutionary responses to epidemics. In comparison to the mixed-effects analytical models from Q1 and Q2, the categorical moderator for splitting host-parasite (co)-evolution into three separate parts was dropped and the host-parasite (co)-evolution was analysed collectively. This approach overcame the challenge of performing comparative analysis across a large number of moderator levels and concurrent low sample sizes. Similarly, in contrast to the mixed-effects meta-analytical model that included a three-way interaction term (Q2), a set of three models were developed that each included only a single categorical moderator according to the specific hypothesis under investigation (Table 3). The models testing the individual moderators were fitted to the reduced dataset excluding the extremely large effect size (lnRR = 4.14, Auld et al., 2014) because of a greater risk of influencing the overall effects of moderators with multiple levels and a low number of total effect sizes / studies.

One model required an alternative optimisation routine (nlm) because the default optimiser did not achieve convergence. As the optimiser determines the numerical algorithm used to estimate model parameters rather than the statistical specification of the model, this adjustment was not expected to influence comparisons among models. Convergence was assessed after refitting, and parameter estimates were retained from the converged solution.

As previously explained under the fourth point of the *Data collection* section (entitled *Processing*), only a single study measured non-additive host–parasite coevolution (Paplauskas et al., 2021), and its data could not be used to calculate lnRR effect sizes. Therefore, during the individual moderator analysis, there was no examination of non-additive host–parasite coevolution.

### Sensitivity analysis for the meta-regression model

My overall dataset comprised effect sizes collected from 11 different studies, which only just surpassed the commonly accepted total of 10 studies for meta-analysis below which there are potential statistical biases…(references). To be completely rigorous, I performed three types of sensitivity analysis including (i) adapting models to explore heterogeneity and the effects of different variables (see Q2 and Q3), (ii) refitting meta-analytical models to a reduced version of the data that excluded a potential outlier (see Q1-3) and (iii) performing two ‘leave-one-out’ analyses for the results of the meta-regression model (Higgins et al., 2024). Leave-one-out analyses were performed for the meta-regression model due to their greater complexity than the corresponding meta-analyses that potentially made them more sensitive to the influence of individual studies or datapoints. This latter approach is described in the following.

Two sensitivity analyses were performed for the meta-regression model using the ‘leave-one-out’ method (Higgins et al., 2024). The first involved the iterative exclusion of one study at a time, separately for each component of host–parasite (co)-evolution × host-class combination for which more than one study was available. Because only one study was available for each of the plant and bacterial host classes, these analyses were restricted to the invertebrate host class and were performed for the three components of host–parasite (co)-evolution represented in the dataset: host evolution, parasite evolution, and net coevolution. The second series involved the iterative exclusion of one independent treatment–control pair (replicate) at a time from the data used to calculate effect sizes, thereby assessing the sensitivity of the meta-regression results to individual replicate pairs contributing to the effect-size estimates. The pre-identified possible outlier was not removed from either of these sensitivity analyses (lnRR = 4.14; Auld et al., 2014), as they were intended to assess the robustness of results to the influence of potentially influential observations rather than to remove such observations a priori.

Of the 11 leave-one-study-out models, one failed to converge using the default optimiser when the study by Duffy et al. (2008) was omitted. Because convergence failure alone does not indicate that the omitted study is influential, I refitted this model using an alternative optimiser to obtain a converged solution and enable comparison with the other leave-one-study-out models. Similarly, four of the leave-one-comparison-out models failed to converge using the default optimiser. These models were refitted using the nlm optimiser to obtain converged solutions and enable comparison with the other leave-one-comparison-out models. As with the leave-one-study-out analyses, convergence failure was not interpreted as evidence that the omitted comparison was influential.

## Results

### Invertebrate (*Daphnia*) host-parasite populations dominate my dataset

My overall dataset contained a combination of effect 123 sizes from 11 different studies corresponding to three components of host-parasite (co)-evolution (host evolution, parasite evolution and net coevolution). In parallel, I analysed a small sample of non-effect size data attributable to 16 cases of non-additive *Daphnia* (invertebrate) host *–* bacterial microparasite coevolution that were derived from one of the studies included in the overall dataset. Similar to the sample of non-effect size data, the overall dataset was heavily biased towards studies of invertebrate host-parasite systems (9 out of 11 studies; Fig. 2). These were mainly representative of *Daphnia magna* and *Daphnia dentifera* in association with either the highly virulent sterilising bacterial parasite *Pasteuria ramosa* or the less intensively studied fungal yeast parasite *Metschinkowa bicuspidata* (7 and 2 out of 9 studies respectively; Fig. 2). There were only two studies representative of non-invertebrate host-parasite systems, including a pair of plant (*Linum usitatissimum*) and bacterial (*Pseudomonas fluorescens*) host-parasite systems (Fig. 2).

**Figure 2.**
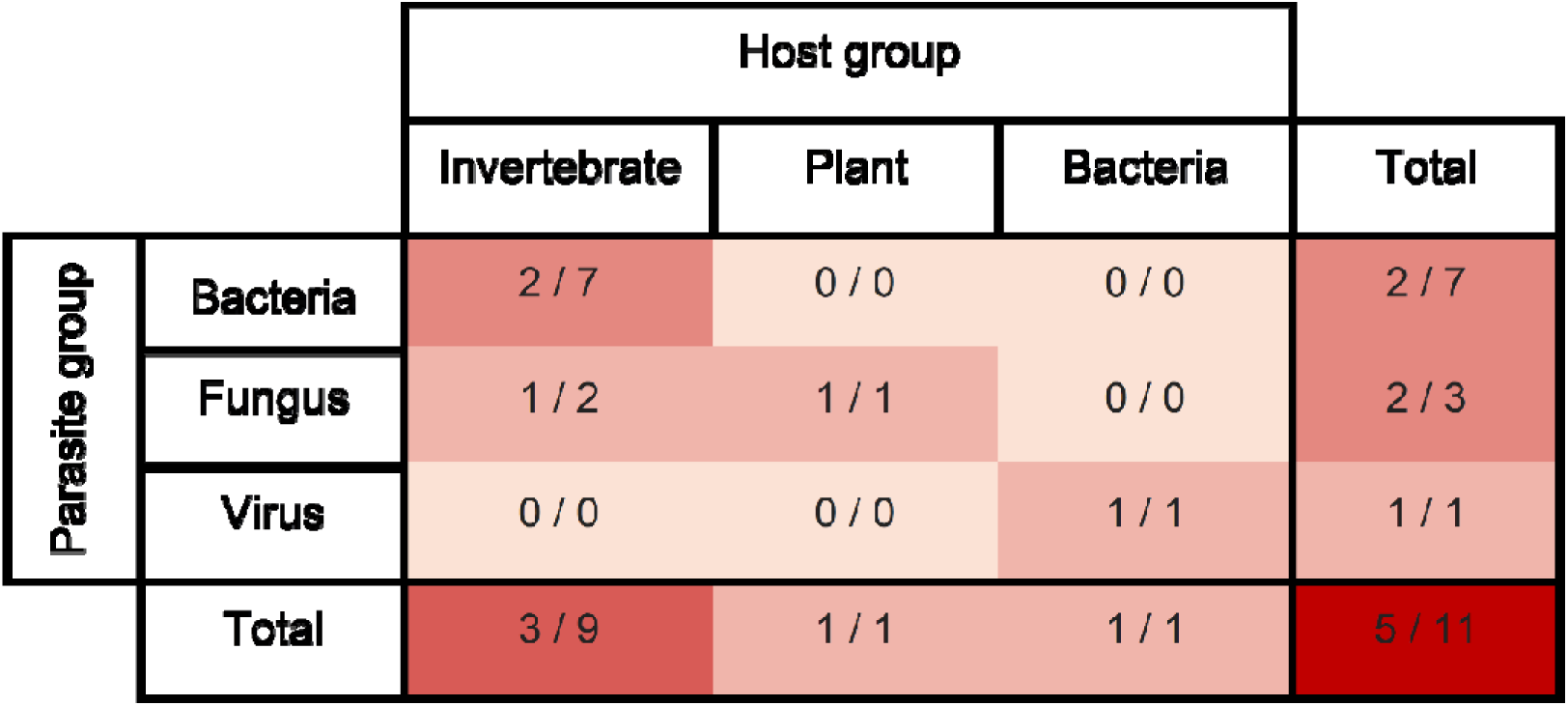
The paired distribution of unique host and parasite species within the data. To avoid conflating the apparent breadth of host and parasite taxa within the dataset by presenting them in separate tables, the number of unique combinations of host and parasite species are shown in each cell of the table, along with the number of studies they are sourced from after a backslash. The colour system corresponds to the number of unique combinations of host and parasite genera, where higher numbers have darker colouration.

Despite their limited representation at the study-level, the representation of plant and bacterial host-parasite systems at the effect size-level was relatively high. This included 24 out of 56 effects sizes for host evolution (evenly distributed between plant and bacterial host-parasite systems) and 12 out of 49 effect sizes for net coevolution (exclusively plant host-parasite systems (later highlighted in Fig. 4). Therefore, plant and bacterial host-parasite systems are relatively well-represented at the effect size-level. However, a notable limitation of the data is that there is substantial variation in the number of effect sizes corresponding to different host-parasite systems across different components of host-parasite (co)-evolution (see later Fig. 3 and 4). This could drastically limit my ability to generalise about any possible relationships between epidemics (and their size) and different components of host-parasite (co)-evolution.

**Figure 3.**
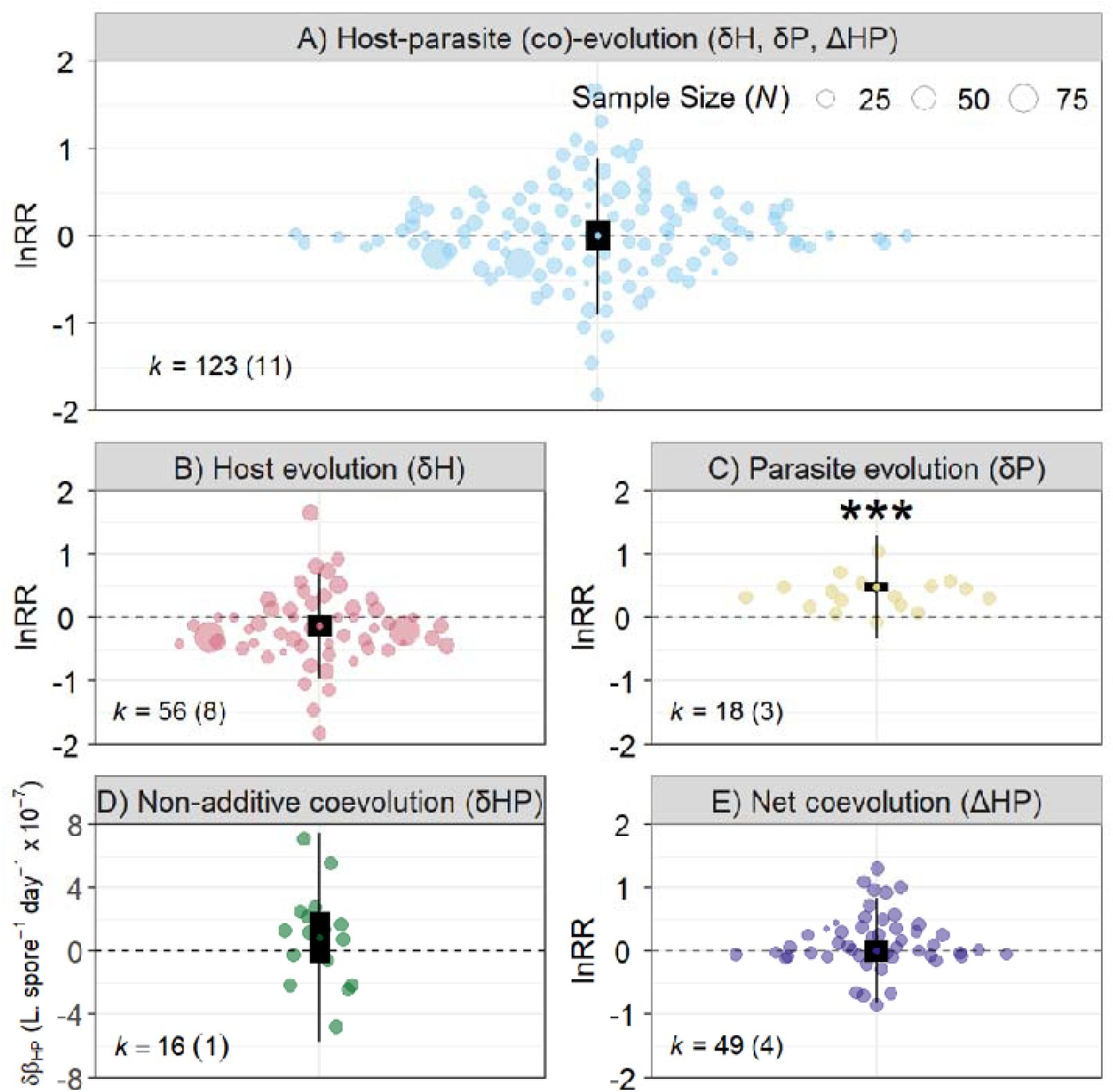
Patterns of host-parasite (co)-evolution that generally occur as a result of epidemics. Effect size data was measured in terms of the log-response ratio (lnRR) for three out of four components of host-parasite (co)-evolution (host evolution [δH], parasite evolution [δP] and net host-parasite coevolution [ΔHP], either collectively (A), or individually (B, C, E), alongside non-effect size values for change in parasite transmission rate owing to non-additive coevolution (δHP, D). Note that since host and parasites naturally share infection as a phenotype, despite different components of host-parasite (co)-evolution being measured on the same scale, positive values of lnRR correspond with phenotypic changes promoting infection, whereas negative values of lnRR correspond to the opposite pattern. The magnitude and direction of effect sizes is relative to lnRR = 0 (horizontal dashed line), which represents no change between the means of pre- and post-epidemic population-level phenotypes. Group-level mean effect sizes are highlighted in black (thick and thin lines are representative of confidence and prediction intervals). The significance of the weighted group-level mean effect for the evolution of parasite infectivity is indicated by the asterisks in bold font (p < 0.001). Effects sizes are scaled according to sample size. The number of replicate host or parasite phenotypes used for calculating average phenotypes within populations at each epidemic timepoint (i.e. sample size) was used to indicate effect size weighting (N). *k* = number of effect sizes (number of studies). Note the number of effects includes the outlier value for parasite evolution (lnRR = 4.14; Auld et al., 2014) which sits outside of the y-axis scale (optimised for readability).

**Figure 4.**
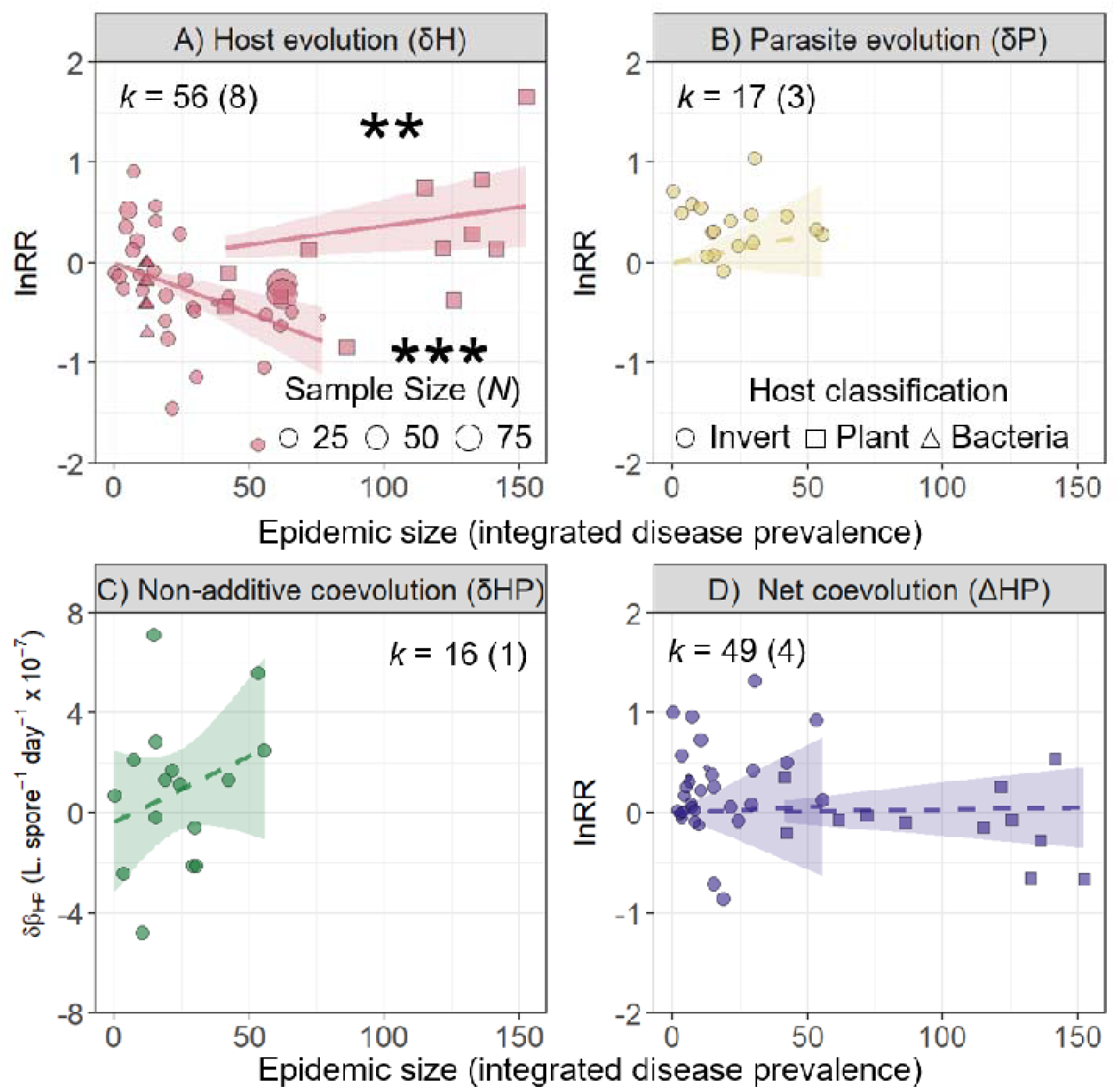
The relationship between epidemic size and different components of host-parasite (co)-evolution. Excluding the collective analysis of three out of four components of host-parasite (co)-evolution from Fig. 3, but consistent with its detailed breakdown, the relationship between epidemic size and four different components of host-parasite (co)-evolution is shown for three host classifications, including invertebrates (circle), plants (square) and bacteria (triangle). Effect size data was measured in terms of the log-response ratio (lnRR) for three out of four components of host-parasite (co)-evolution, including A) host evolution (δH), B) parasite evolution (δP) and D) net host-parasite coevolution (ΔHP), alongside non-effect size values for change in parasite transmission rate owing to non-additive coevolution (δHP, D). Note that since host and parasites naturally share infection as a phenotype, despite different components of host-parasite (co)-evolution being measured on the same scale, positive values of lnRR correspond with phenotypic changes promoting infection, whereas negative values of lnRR correspond to the opposite pattern. The magnitude and direction of effect sizes is relative to lnRR = 0 (horizontal dashed line), which represents no change between the means of pre-and post-epidemic population-level phenotypes. Effects sizes are scaled according to sample size. The number of replicate host or parasite phenotypes used for calculating average phenotypes within populations at each epidemic timepoint (i.e. sample size) was used to indicate effect size weighting (N). Lines represent the meta-regression coefficients for each host classification (excluding bacteria for greater readability), where solid versus dashed lines refer to significant (p < 0.05) and non-significant estimates respectively. The shaded bands correspond to the 95% confidence intervals for each estimate. The number of asterisks indicate significance thresholds (two and three for p < 0.01 and p < 0.001 respectively). *k* = number of effect sizes (number of studies). Note the number of effects includes the outlier value for parasite evolution (lnRR = 4.14; Auld et al., 2014) which sits outside of the y-axis scale (optimised for readability).

### Epidemics drive the evolution of greater parasite infectivity

To characterise general host-parasite (co)-evolutionary responses to epidemics, the overall pattern for three components of host-parasite (co)-evolution (host evolution, parasite evolution and net coevolution) was quantified prior to a detailed breakdown of its component parts (including non-effect size data for non-additive coevolution; Fig. 3).

There was no overall pattern in how host-parasite (co)-evolution generally occurs as a result of epidemics when analysing three components of (co)-evolution simultaneously (lnRR = - 0.01, 95% CI = [-0.17, 0.17], p = 0.99; Fig. 3A). In comparison, when comparing amongst individual components of host-parasite (co)-evolution, whilst there were no general patterns of host evolution (lnRR = -0.13, 95% CI = [-0.29, 0.35], p = 0.12; Fig. 3B), non-additive coevolution (one-sample t-test: M = 8.57 × 10⁻□, t₁₅ = 1.15, 95% CI [−7.38 × 10⁻□, 2.45 × 10⁻□], p = 0.27); Fig. 3C) and net coevolution (lnRR = 0.01, 95% CI = [-0.18, 0.18], p = 0.97; Fig. 3D), there was a very consistent pattern of parasite evolution (lnRR = 0.49, 95% CI = [0.41, 0.56], p < 0.001; Fig. 3B). This same result was still significant a putative outlier was removed (lnRR = 0.48, 95% CI = [0.42, 0.54], p < 0.001). This strongly suggests that epidemics, regardless of size, very consistently drive parasite populations towards the evolution of greater infectivity.

Heterogeneity analysis for the individual breakdown of host-parasite (co)-evolutionary responses that generally occur as a result of epidemics showed that the total variation explained by between-study differences was very high (I^2^ > 75%). Although a large proportion of this heterogeneity was explained by the random effects for host and parasite genus (31.4% and 15.0% respectively), there was a relatively large remaining proportion of unexplained heterogeneity. In addition, taking into account the highly significant result for parasite evolution and the consistency in the direction of individual effect sizes, this high level of heterogeneity was likely driven by the host evolution and net coevolution components of host-parasite coevolution. The sources of heterogeneity for these components of host-parasite coevolution was examined in subsequent analyses.

### Larger epidemics drive greater host evolution

Building on my previous analysis that involved characterising general host-parasite (co)-evolutionary responses to epidemics, regardless of size, I quantified the relationship between epidemic size and the strength of four different components of host-parasite (co)-evolution. This involved assigning host-parasite systems into three taxonomic groups based on a broad classification of hosts as either invertebrates, plants or bacteria, and introducing a continuous moderator variable for epidemic size into my earlier meta-analytical model to perform a meta-regression. This also allowed me to identify contributing factors to the high heterogeneity level observed as part of the previous analysis.

In contrast to the absence of any general pattern of host evolution in response to epidemics (Fig. 3B), which could also have been a large contributing factor to the aforementioned high level of heterogeneity observed as part of the previous analysis, the strength of invertebrate and plant host evolution was strongly determined by epidemic size (β = -0.010, 95% CI = [-0.015, -0.006], p < 0.0001 and β = 0.004, 95% CI = [0.001, 0.006], p = 0.006 respectively; Fig. 4A). Unexpectedly, the direction of host evolution varied between invertebrate and plant host classifications, such that larger epidemics drove the evolution of reduced susceptibility to infection for invertebrate hosts, whereas they drove increased susceptibility for plant hosts. Despite the lack of any significant effect of epidemic size on bacterial hosts’ evolution, the estimated relationship was negative (β = -0.019, 95% CI = [-0.048, 0.010], p = 0.188). Note that the meta-regression line was removed from Fig. 4A due to limited epidemic-size categories (11.7, 12.2), reducing the independent variation available to estimate the meta-regression slope.

Again, in contrast to the general pattern of the evolution of parasites towards greater infectivity in response to epidemics (Fig. 3C), there was no relationship between epidemic size and parasite evolution (β = 0.006, 95% CI = [-0.003, 0.014], p = 0.187; Fig. 4B). This result was unchanged when the outlier was removed (β = 0.003, 95% CI = [-0.07, 0.013], p = 0.556; Fig. 4B). Despite the estimated relationship being positive, this was not significant. Similar to the result for parasite evolution, the positive estimated relationship between epidemic size and non-additive host-parasite coevolution was not significant (estimate = 5.27 × 10⁻□ ± 4.71 × 10⁻□ SE, t₁₄ = 1.12, p = 0.281, R^2^ = 0.082; Fig. 4C). Finally, there was no relationship between epidemic size and net coevolution (β = 0.002, 95% CI = [-0.012, 0.015], p = 0.794 and β = 0.001, 95% CI = [-0.002, 0.004], p = 0.631 for invertebrate and plant host-parasite systems respectively; Fig. 4C).

### Contextual influences on host–parasite (co)-evolution

In addition to characterising patterns of host–parasite (co)-evolution that occur during epidemics and their relationship with epidemic size, I used moderator analyses to test hypotheses concerning the potential influence of three contextual factors on host–parasite (co)-evolution (Table 3). These analyses considered three of the four components of host–parasite (co)-evolution represented in my dataset in combination: host evolution, parasite evolution, and net coevolution.

Moderator analysis indicated that the measure used to quantify change in host–parasite (co)-evolution influenced effect sizes. Estimates were close to zero for transmission rate (lnRR = 0.134, 95% CI = −0.153 to 0.422, p = 0.360; Fig. 5A) and prevalence of infected hosts (lnRR = 0.008, 95% CI = −0.294 to 0.309, p = 0.961; Fig. 5A). In contrast, prevalence of susceptotypes showed a significant negative effect (lnRR = −0.548, 95% CI = −0.944 to −0.152, p = 0.007; Fig. 5A), whereas the plaque-forming-unit-based measure was also negative but not statistically significant (lnRR = −0.191, 95% CI = −0.641 to 0.259, p = 0.405; Fig. 5A).

**Figure 5.**
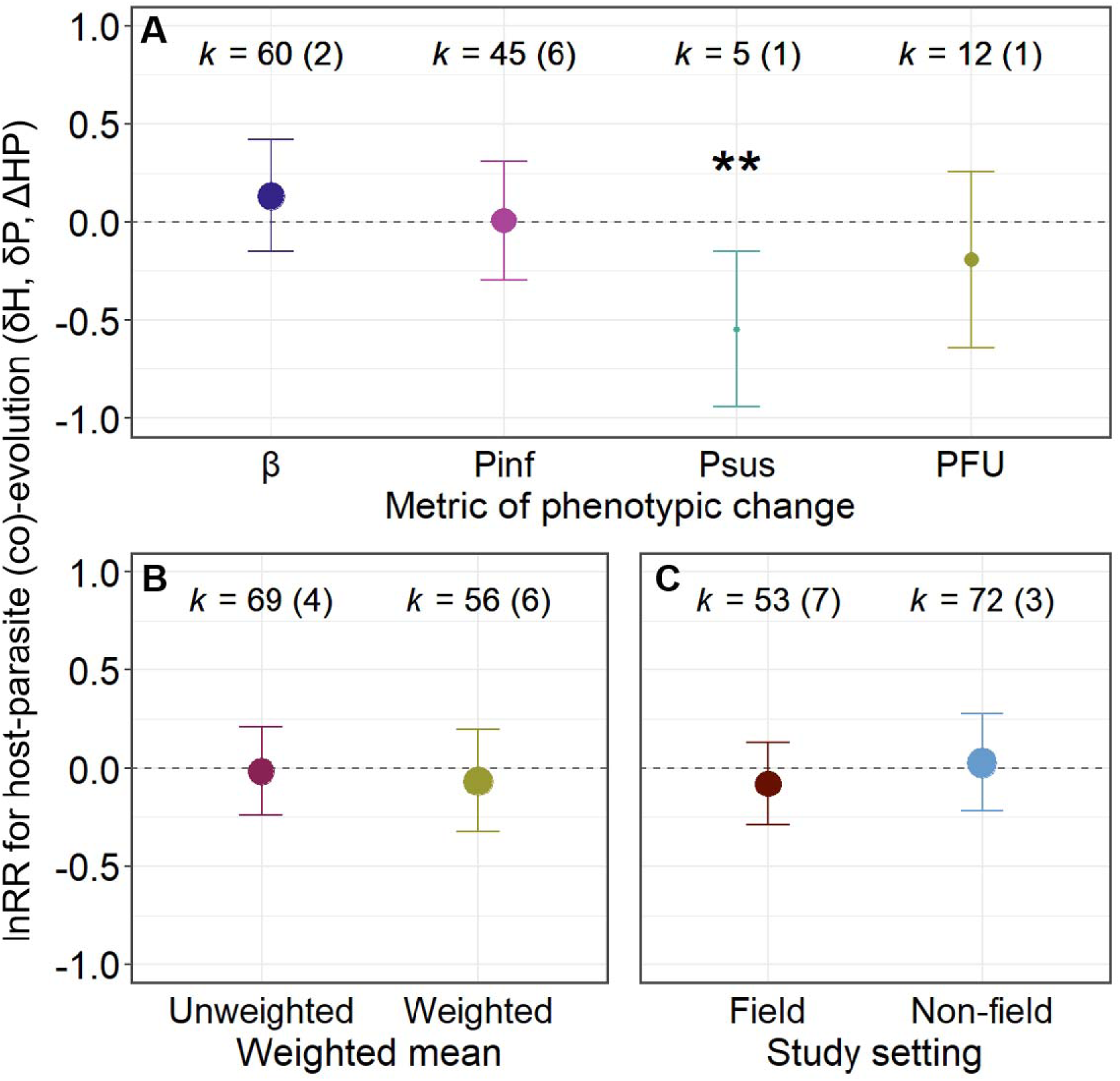
Wider factors that influence host-parasite (co)-evolutionary responses to epidemics. These analyses considered three of the four components of host–parasite (co)-evolution represented in my dataset in combination: host evolution (δH), parasite evolution (δP) and net host-parasite coevolution (ΔHP). Effect sizes were measured as log-response ratios (lnRR). A) Metric of phenotypic change: (i) proportion of infected individuals within a host population (Pinf), (ii) parasite transmission rate (β), (iii) proportion of host clones within a population susceptible to parasite attachment (Psus; as opposed to resistotypes *sensu* Ameline et al., 2022), and (iv) proportion of plaque-forming units on bacterial colonies (PFU; Gomez and Buckling, 2011). B) Comparison of frequency-weighted and unweighted means. C) Study setting (field versus non-field). Because host and parasite responses ultimately share infection as a common phenotype, positive lnRR values indicate phenotypic changes that promote infection, whereas negative values indicate changes in the opposite direction. Effect sizes are interpreted relative to lnRR = 0 (horizontal dashed line), which represents no change between pre- and post-epidemic population-level phenotypes. Unlike Figures 3 and 4, which show individual effects, this figure presents overall model coefficients; accordingly, datapoint sizes are scaled by the number of effects (rather than sample size). *k* denotes the number of effects, with the number of studies given in parentheses. ** indicates *p* < 0.01.

Interpretation of the moderator effect of how change in host–parasite (co)-evolution was measured should be treated cautiously because the representation of moderator levels was highly unbalanced. Pinf and β were represented by 60 and 45 effect sizes from 2 and 6 studies, respectively, whereas prevalence of susceptotypes and PFU were represented by only 5 and 12 effect sizes, respectively, both from a single study. Thus, the more pronounced negative estimates for prevalence of susceptotypes and PFU may reflect study-specific effects or differences in the biological contexts represented by these measures, rather than systematic differences among response variables. In addition, differences in host classification may represent an unaccounted source of heterogeneity underlying the apparent moderator effect, particularly if host classifications were unevenly represented across moderator levels.

I also tested whether methodological characteristics of the studies influenced estimates of change in host–parasite (co)-evolution. Specifically, I assessed whether the weighting approach used to calculate effect sizes and whether studies were based on field or non-field sampling influenced the estimated change. Neither weighting approach was associated with a significant effect, with the weighted approach showing a negative estimate (lnRR = −0.064, 95% CI = −0.327 to 0.199, p = 0.634; Fig. 5B) and the unweighted approach showing a near-zero estimate (lnRR = −0.016, 95% CI = −0.242 to 0.211, p = 0.892; Fig. 5B). Similarly, field sampling status was not associated with a significant effect, with field studies showing a negative estimate (lnRR = −0.080, 95% CI = −0.290 to 0.130, p = 0.456; Fig. 5C) and non-field studies showing a near-zero positive estimate (lnRR = 0.028, 95% CI = −0.221 to 0.277, p = 0.827; Fig. 5C).

### Meta-regression results were robust to data exclusion

Two leave-one-out sensitivity analyses were performed for the meta-regression, involving the iterative exclusion of either one study or one independent treatment–control pair (replicate) used to calculate an effect size at a time (Table 1; Fig. S2 and S3). Overall, the direction of estimated relationships between host–parasite (co)-evolution and epidemic size was consistent with those of the original meta-regression, although the magnitude and statistical significance of some relationships varied among leave-one-out iterations and between study- and replicate-level analyses.

At the study level, the relationship between epidemic size and host evolution for invertebrate host–parasite systems remained negative (estimate = −0.009 ± 0.001, SE = 0.002 ± 0, p < 0.001). The relationship between epidemic size and parasite evolution was positive but non-significant (estimate = 0.026 ± 0.021, SE = 0.010 ± 0.005, p = 0.203 ± 0.177). Similarly, the relationship between epidemic size and net coevolution remained positive but non-significant (estimate = 0.003 ± 0.001, SE = 0.006 ± 0.001, p = 0.630 ± 0.136).

At the replicate level, the negative relationship between epidemic size and host evolution for invertebrate systems remained significant (estimate = −0.010 ± 0, SE = 0.002 ± 0, p < 0.001), as did the positive relationship for plant systems (estimate = 0.004 ± 0, SE = 0.001 ± 0, p = 0.020 ± 0.012). The relationship between epidemic size and host evolution for bacterial systems was negative but non-significant (estimate = −0.019 ± 0.001, SE = 0.010 ± 0.002, p = 0.171 ± 0.043). Relationships between epidemic size and parasite evolution in invertebrate systems (estimate = 0.006 ± 0, SE = 0.004 ± 0, p = 0.168 ± 0.024), net coevolution in invertebrate systems (estimate = 0.001 ± 0, SE = 0.006 ± 0, p = 0.845 ± 0.012), and net coevolution in plant systems (estimate = 0 ± 0, SE = 0.001 ± 0, p = 0.751 ± 0.037) were also non-significant.

Overall, the sensitivity analyses indicated that the estimated relationships were generally consistent in direction with the original meta-regression, although statistical significance depended on whether studies or individual replicate pairs were excluded. The consistency of the negative association between epidemic size and host evolution in invertebrate systems across both sensitivity analyses provides evidence that this relationship was not driven by a single study or replicate pair.

## Discussion

The intentions behind this study were two-fold: to characterise general relationships between epidemic size and different forms of host–parasite (co)-evolution, and to overcome the lack of replication in a literature heavily biased towards *Daphnia* host–parasite systems. However, despite targeting a broader literature, the final dataset remained strongly biased towards *Daphnia*, comprising just 123 effect sizes from 11 studies that rigorously integrated measures of epidemic size and (co)-evolution. Across this limited evidence base, host evolutionary responses and net coevolutionary change varied substantially in direction, whereas parasite infectivity showed a more consistent increase. Epidemic size partly explained this variation in host evolution, with larger epidemics associated with reduced susceptibility in invertebrates but increased susceptibility in plants, while parasite evolution, non-additive coevolution and net coevolution were largely independent of epidemic size. Thus, the results provided partial support for my hypotheses: the predicted association between larger epidemics and reduced host susceptibility was supported for invertebrates, but not plants, whereas the predicted independence of parasite evolution from epidemic size was supported.

The association between epidemic size and host evolutionary change in invertebrates provides support for the original hypothesis that larger epidemics impose stronger selection for reduced host susceptibility. There are nevertheless some nuances to this general pattern, as the evolutionary response to epidemic size may depend on the relative importance of epidemic-driven selection compared with other biotic and abiotic pressures. Among the studies included in this meta-analysis, Duffy et al. (2012) found that the direction of susceptibility evolution in *Daphnia* depended on the combination of epidemic size and host productivity. Strauss et al. (2017) showed that parasite-driven evolution of host competitive ability could increase host density and consequently support larger epidemics, while Walsman et al. (2023) demonstrated that sufficiently high parasite abundance can favour lower resistance when its costs outweigh its benefits. Together, these findings support a general association between larger epidemics and selection on host susceptibility, while highlighting that the evolutionary response can also depend on the broader selective environment.

Relationships with epidemic size may also differ among components of host and parasite fitness and over different timescales. The present study focused primarily on transmission-related traits, whereas virulence-related traits can show different patterns. Among the studies included in this meta-analysis, Auld and Brand (2017) found that parasite spore production increased independently of epidemic size, while host susceptibility to parasite virulence increased with epidemic size. Similarly, Gowler et al. (2023) found no change in parasite virulence over an epidemic but a reduction in spore production, which they suggested could reflect trade-offs associated with parasite growth. Ameline et al. (2022) further showed that genetic slippage can cause evolutionary changes generated during an epidemic to have limited consequences for longer-term evolutionary trajectories. Thus, evolutionary responses to epidemics varied among fitness-related traits and may not persist beyond the epidemic itself.

Host and parasite generation times are an important potential confounding factor in cross-system comparisons, as they determine the temporal scale over which evolutionary responses to epidemics can occur. In plants, for example, longer host generation times may limit evolutionary responses to epidemic-driven selection (Brockhurst and Koskella, 2013), while frequency-dependent selection (Thrall et al., 2012) and sparse sampling of epidemic size may also contribute to the contrasting plant result (Thrall et al., 2012, personal communication). In bacteria, the estimated relationship was strongly negative and consistent with our prediction, but was not statistically significant. Ideally, differences in host and parasite generation times would be incorporated when comparing epidemic size across systems, but reliable estimates are not available for all host–parasite combinations represented here. Together, these factors suggest that variation among host groups may reflect differences in the temporal and ecological context of evolution, rather than a general tendency for larger epidemics to produce stronger evolutionary responses.

For bacteria, the estimated relationship was negative, consistent with our prediction, but non-significant. However, interpretation is limited by the restricted variation in epidemic size, with estimates available only as study-level means and falling into two categories (11.7 and 12.2). Moreover, epidemic size was estimated as PFU/(CFU + PFU), an adapted measure of infection prevalence that may not directly reflect the proportion of infected host cells. In the original *Pseudomonas fluorescens*–phage system, Gómez and Buckling (2011) quantified bacterial and phage densities separately through time, capturing both host abundance and phage exposure, and observed fluctuating selection dynamics. Using PFU alone as an alternative proxy for pathogen abundance or exposure produced a similar pattern in our analysis, but measures of infected-cell prevalence or density, particularly when tracked through time, may provide a more direct measure of epidemic size. This distinction may be important because PFU reflects infectious phage particles rather than infected bacterial cells, and phage abundance can vary independently of host infection prevalence through differences in phage production and persistence.

The moderator analyses provide some indication that the apparent magnitude of host–parasite (co)-evolution depends on how evolutionary change is quantified. Measures more closely related to host susceptibility, particularly prevalence of susceptotypes, showed more pronounced negative estimates than measures of realised infection or transmission, which were close to zero. This pattern should be interpreted cautiously because the susceptibility-related measures were represented by only one study each, and therefore may reflect study-specific biological or methodological characteristics rather than a general effect of measurement type. Differences in host classification may also contribute to this heterogeneity if particular measures were preferentially applied to particular host–parasite systems. Nevertheless, the concentration of negative estimates for measures more directly characterising host susceptibility suggests that effects may become more apparent when assessed at the level of host susceptibility rather than realised population-level infection, although this interpretation requires confirmation from a more balanced dataset.

In contrast, there was little evidence that the methodological characteristics examined here systematically influenced estimates of evolutionary change. Neither the use of frequency weighting nor whether sampling occurred in the field was associated with significant differences in effect sizes. The lack of an effect of frequency weighting suggests that the overall patterns were not strongly dependent on whether common genotypes contributed more to population-level estimates. However, frequency weighting necessarily emphasises the evolutionary responses of common genotypes and may therefore obscure variation among less frequent genotypes, which could be important for understanding the consistency of evolutionary responses within populations. Thus, while weighting appears unlikely to explain the broad patterns observed here, examining variation among genotypes and replicate populations may provide additional insight into the consistency of host–parasite (co)-evolution.

Beyond the effects of epidemic size, the variability of host–parasite (co)-evolution may depend on asymmetry in host and parasite evolutionary rates. Parasites are generally expected to evolve more rapidly and consistently than hosts because of their larger population sizes and shorter generation times (Schmid-Hempel, 2011), potentially causing hosts to lag behind parasite adaptation. The greater consistency of parasite infectivity than host evolution observed here is consistent with this expectation, although it does not indicate how often either antagonist is locally adapted or maladapted. This could be examined using the local-adaptation framework by testing whether asymmetry in evolutionary rates is associated with the frequency or magnitude of local adaptation. One approach would be to reanalyse the data underlying previous local-adaptation meta-analyses (Greischar & Koskella, 2007; Hereford, 2009; Hoeksema & Forde, 2009), where available, using measures of variability such as lnCVR to test whether hosts or parasites show greater consistency in fitness across environments. This would complement the present focus on the magnitude and direction of evolutionary change by examining how consistently reciprocal adaptation occurs.

Two additional sources of inconsistency concern the terminology and measurement of epidemic size (Table S2). Studies frequently use terms such as disease prevalence, infection prevalence and parasite prevalence interchangeably, despite these measures not necessarily being equivalent. Methods for quantifying prevalence also vary considerably, including sampling intensity and criteria for confirming infection, while definitions of an “epidemic” are rarely stated explicitly. These inconsistencies may limit the comparability of epidemic-size estimates and therefore the interpretation of relationships with host–parasite (co)-evolution.

Overall, this synthesis provides limited evidence that epidemic size consistently determines the magnitude or direction of host–parasite (co)-evolution. Larger epidemics were associated with greater host evolutionary change in invertebrates, consistent with stronger selection for reduced susceptibility, but this relationship was reversed in plants and non-significant in bacteria. In contrast, parasite evolution, non-additive coevolution and net coevolution showed no detectable relationship with epidemic size, despite a relatively consistent increase in parasite infectivity across studies. These contrasting patterns highlight the importance of host and parasite generation times, the relative strength of epidemic-driven selection compared with other biotic and abiotic pressures, and the ecological and temporal context in which evolution occurs. Interpretation is further constrained by substantial heterogeneity in how epidemic size and evolutionary change were quantified, including inconsistent definitions of infection, disease and epidemics. The strong representation of *Daphnia* systems and the small number of studies also limit the generality of the conclusions.

A major priority is to address the limited empirical evidence linking epidemic dynamics with host–parasite (co)-evolution. More studies should track evolutionary change and disease prevalence through time, particularly using experimental time-shift approaches that directly assess reciprocal evolutionary responses (Brockhurst and Koskella, 2013). Broadening inclusion beyond time-shift experiments to other measures of coevolution could help expand the evidence base, although this may come at the cost of less direct evidence for reciprocal evolutionary change. This work also represents an empirical step towards linking the Disease Cycle framework with quantitative epidemiological models. As proposed by Paplauskas et al. (2025), relationships between epidemic size and host evolution, parasite evolution and net coevolution could be incorporated into existing epidemiological models without explicitly modelling the underlying coevolutionary mechanisms. Future work should therefore seek to generate robust quantitative estimates that can be used to incorporate coevolutionary feedbacks into models of disease dynamics.

## Author contributions

Sam Paplauskas: conceptualization (lead), formal analysis (lead), investigation (lead), methodology (lead), project administration (lead), visualization (lead), writing – original draft (lead), writing – review and editing (lead).

## Acknowledgements

Thanks go to my former PhD supervisor, Dr Stuart Auld, for their support during the initial stages of the research. Thanks go to Assistant Professor Alexander Strauss, Dr. Camille Ameline, Dr. Peter Thrall and Dr. Jason Walsman for sharing their raw data and clarifying their methods.

## Data availability statement

The data supporting this study are available from the Zenodo repository at https://doi.org/10.5281/zenodo.22828287 (Paplauskas, 2026).

## Supporting information

See the supporting information file.

