## Supplemental Figures and Tables for "Epidemic size and host–parasite (co)-evolution: a meta-analysis"

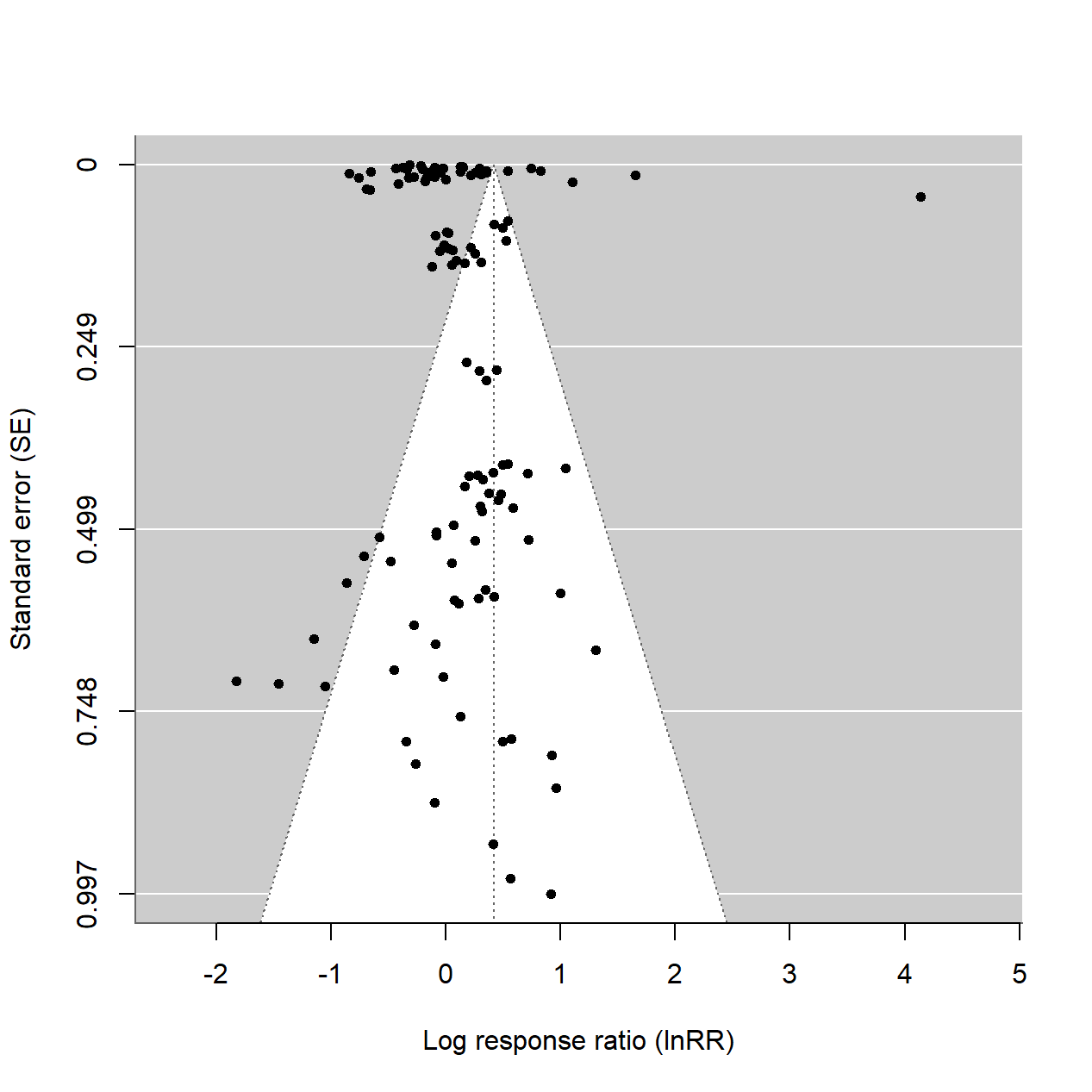

**Supplementary Figure S1. Funnel plot to test for the presence of publication bias.** The distribution of individual log response ratio values around the overall estimate from a mixed-effects meta-analytical model (dashed lines show both the overall model estimate and 95% confidence intervals) according to their standard error (SE).

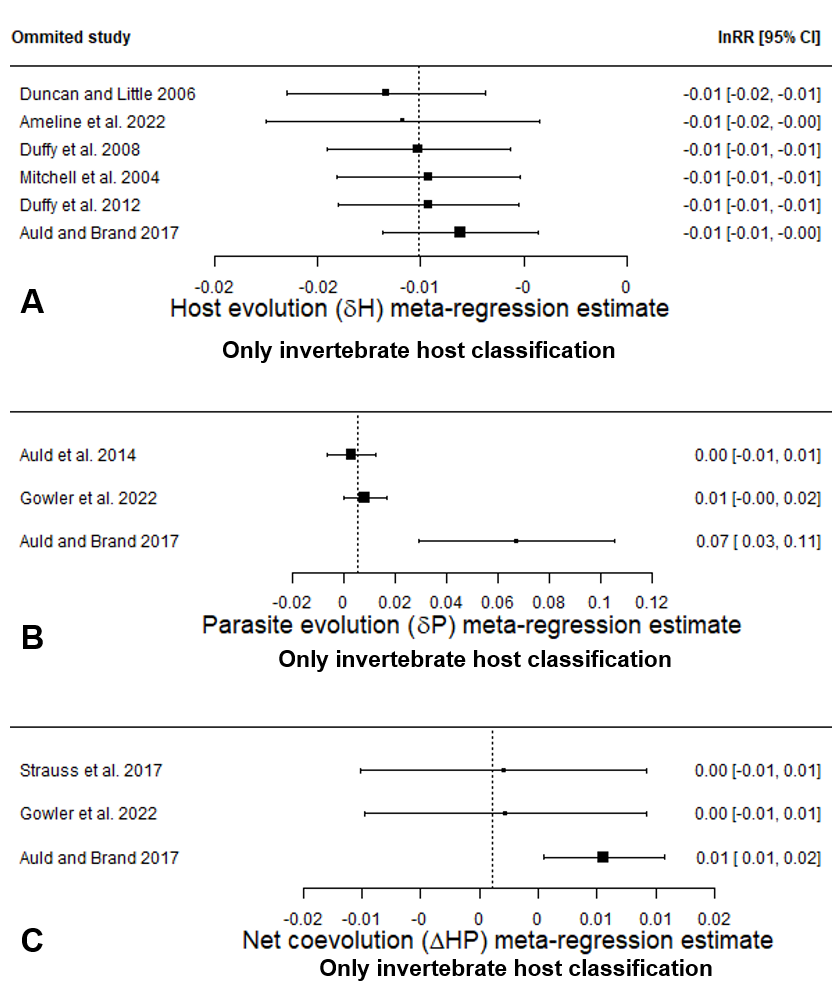

**Supplementary Figure S2. Forest plots corresponding to meta-regression estimates for the leave one study out sensitivity analysis for only invertebrate host-parasite systems.** The meta-regression represents a three-way interaction term between different components of host-parasite (co)-evolution (A-C), host taxonomic classification and epidemic size. Leave-one-out sensitivity analyses are shown for invertebrate host–parasite systems only; plant and bacterial systems, each represented by a single study, are not shown. Iterative exclusion of these studies was also omitted because removing a single study from these sparsely represented host groups was not expected to substantially alter the overall model estimates. Each datapoint corresponds to the estimate from a different iteration of a meta-regression model using the same structure, where a different study has been omitted (authors and values on the left and right hand-side margins of the plot respectively). In contrast to a traditional forest plot, where the size of each datapoint is scaled according to its relative contribution to the overall mean, the size of each datapoint in these graphs indicates the precision of each model (1 / SE). The bars represent 95% confidence intervals (square brackets on the right-hand side) and the meta-regression estimate of the original model using the full set of studies is shown by the dotted line. Note that a pre-identified possible outlier has not been removed for sensitivity analysis (lnRR = 4.14; Auld et al., 2014), as sensitivity analysis is specifically intended to assess the robustness of results to the influence of potentially influential observations rather than to remove such observations a priori.

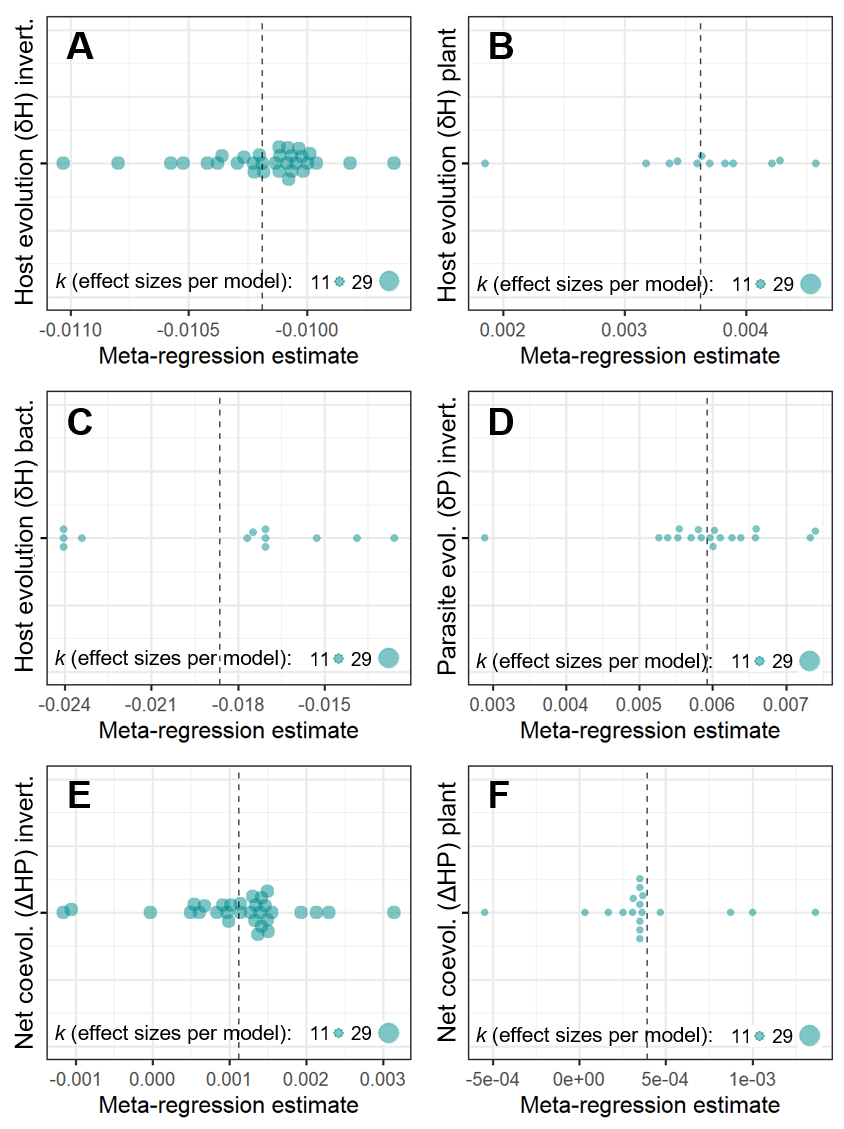

**Supplementary Figure S3. Adapted orchard plots corresponding to meta-regression estimates for the leave one independent comparison out sensitivity analysis across all host-parasite system classifications.** The meta-regression represents a three-way interaction term between three out of four components of host-parasite (co)-evolution (host evolution A-C, parasite evolution D and net coevolution E-F), host taxonomic classification (invertebrates A,C and E, plants B and F and bacteria C) and epidemic size. Each datapoint corresponds to the estimate from a different iteration of a meta-regression model using the same structure, where a different comparison has been omitted. Similar to the difference between Supplementary Figure S2 and traditional forest plots, unlike a traditional orchard plot, where the size of each datapoint is scaled according to its relative contribution to the overall mean, the size of each datapoint in these graphs indicates the number of effect sizes per model iteration. This is indicative of the level of error for each model iteration (equivalent to the number of model iterations per group). The meta-regression estimate of the original model using the full set of studies is shown by the dotted line. Note that a pre-identified possible outlier has not been removed for sensitivity analysis (lnRR = 4.14; Auld et al., 2014), as sensitivity analysis is specifically intended to assess the robustness of results to the influence of potentially influential observations rather than to remove such observations a priori.

Table S1. Google Scholar (GS) search terms listed in descending order according to their number of matches. In comparison to the first search term in the table, only search terms that only appear exclusively in the title can be specified by using the ‘allintitle:’ prefix. In addition, since the GS engine does not follow the traditional Boolean logic, compared to search engines such as PubMed, Web of Science or Scopus, an AND is implicit between every search term separated by a space and a dash (-) can be used to search for an exact phrase.

| **Search term** | **Matches** |
| --- | --- |
| host parasite coevolution epidemic-size | 196 |
| allintitle: antagonistic-coevolution | 129 |
| allintitle: parasite-mediated selection | 85 |
| allintitle: host-parasite epidemic | 13 |
| allintitle: pathogen-epidemic | 13 |

Table S2. Methodological inconsistencies in the measurement and definition of epidemic size across studies included in the meta-analysis.

| **Study** | **Problem type** | **Detailed notes** |
| --- | --- | --- |
| Auld et al. (2014) | Infection-status assessment | Study authors may have applied a subjective approach to verification of infection status, with individuals or samples preferentially selected for dissection to check for signs of infection. |
| Auld et al. (2017) | Sampling intensity | Disease prevalence was calculated from random sampling of the population, but the number of individuals sampled could vary considerably with overall population size. |
| Duffy et al. (2008) | Undefined measure | What “disease prevalence” represented was not clearly specified, making it difficult to determine whether the measure represented visible disease, infection prevalence, or parasite prevalence. |
| Gowler et al. (2022) | Infection-status assessment | Individuals or samples may have been subjectively selected for confirmation of infection status through dissection, potentially introducing inconsistency in how infection was identified. |
| Thrall et al. (2012) | Ambiguous infection definition | It was unclear whether infection status was based on any expression of infection (binary classification) or a threshold level of infection intensity (Ibrahim & Barrett, 1991). |
| Thrall et al. (2012) | Sparse temporal sampling | Integrated epidemic size was estimated from relatively sparse sampling, with as few as two measurements across a season, potentially providing an imprecise representation of cumulative infection and associated selection. |
| Strauss et al. (2017) | Undefined measure | What “disease prevalence” represented was completely omitted, making it unclear what aspect of infection was being quantified. |
| Multiple studies | Terminological inconsistency | Terms such as disease prevalence, infection prevalence and parasite prevalence were used inconsistently. Visible signs of disease cannot necessarily be used to infer infection prevalence or parasite prevalence without direct confirmation of infection. |
| Multiple studies | Undefined epidemic threshold | It was often unclear whether the level of disease measured in each population constituted an “epidemic”, as studies rarely stated an explicit definition despite there being multiple ways to define an epidemic (Paplauskas, 2025). |
